# Evosep One Enables Robust Quantitative Deep Proteome Coverage using Tandem Mass Tags while Significantly Reducing Instrument Time

**DOI:** 10.1101/543371

**Authors:** Jonathan R. Krieger, Leanne E. Wybenga-Groot, Jiefei Tong, Nicolai Bache, Ming S. Tsao, Michael F. Moran

## Abstract

The balance between comprehensively analyzing the proteome and using valuable mass spectrometry time is a genuine challenge in the field of proteomics. Multidimensional fractionation strategies have significantly increased proteome coverage, but often at the cost of increased mass analysis time, despite advances in mass spectrometer acquisition rates. Recently, the Evosep One liquid chromatography system was shown to analyze peptide samples in a high throughput manner without sacrificing in depth proteomics coverage. We demonstrate incorporation of Evosep One technology into our multiplexing workflow for quantitative analysis of tandem mass tag (TMT)-labeled non-small cell lung carcinoma (NSCLC) patient-derived xenografts (PDXs). Using a 30 samples per day Evosep workflow, >12,000 proteins were identified in 48 hours of mass spectrometry time, which is comparable to the number of proteins identified by our conventional concatenated EASY-nLC workflow in 67.5 hours. Shorter Evosep gradient lengths reduced the number of protein identifications by 10%, while decreasing mass analysis time by 50%. Thus, our Evosep workflow enables quantitative analysis of multiplexed samples in less time without conceding depth of proteome coverage.

## Introduction

Comprehensive proteomics has seen an increase in the number of peptides identified and quantified by shotgun strategies due to improved mass spectrometer acquisition rates^1^ and advances in peptide separation strategies. Deep proteome coverage has been achieved by various multidimensional fractionation strategies that involve off- and on-line peptide fractionation methods, which reduce sample complexity prior to tandem mass spectrometry (MS/MS) analysis (reviewed in ^2–6^). The performance of two-dimensional liquid chromatography (2D-LC) depends on the separation efficiency, or peak capacity, in both chromatographic dimensions and orthogonality of the two separation elements ^7^. Fractionation of peptides and proteins increases the peak capacity of LC separation ^8^. Off-line peptide fractionation based on a high pH reversed phase (RP) approach followed by on-line low pH RP provides high resolving power based on hydrophobic interactions, especially when a large number of fractions are collected in the first dimension ^7,9^. However, this increases the number of samples that need to be separated in the second dimension and, consequently, mass spectrometry analysis time. Indeed, high peptide coverage of human cell lines has been observed, but typically at the expense of longer MS acquisition time ^10^. For instance, Nagaraj *et al* identified 10255 human proteins from HeLa cell lysates over 12 days (288 h) using tandem prefractionation by gel filtration at the protein level, digestion by three specific proteases, and strong anion exchange separation of peptides into 72 fractions ^11^. Significant reduction in analysis time has been achieved by concatenating high pH RPLC fractions with minimal reduction (15-20%) in protein identification compared to non-concatenated samples ^12^. Moreover, concatenating multiple early, middle, and late fractions improved the orthogonality of RPLC-RPLC and increased protein sequence coverage when compared to strong cation exchange (SCX)-RPLC schemes ^13^. Using this concatenated high pH RP - low pH RP 2D LC-MS/MS approach, Spicer *et al* achieved deep proteome coverage of a complex sample in relatively little MS analysis time, identifying 8757 proteins from tryptic digests of whole Jurkat cell lysates in 31.5 h on a Triple TOF 5600 instrument (21 fractions) ^4^.

Recently, this concatenated 2D RP-RP LC-MS/MS approach has been applied to quantitative proteomics investigations. The advent of tandem mass spectrometry tags (TMTs) made quantitative proteomics of complex protein samples a reality ^14^. These isobaric labeling reagents ensure that identical peptides labeled with different TMTs exactly comigrate in chromatographic separations, such that peptides from different samples can be accurately quantified ^14^. The desire to profile thousands of proteins from different samples in a quantitative manner, and the need to balance efficiency, statistical power, and throughput led to multiplexing TMT reagents ^15^. Indeed, with TMT 10-plex reagents, up to ten biological samples can be analyzed for proteome changes in a single MS experiment with wide dynamic range and excellent accuracy, on high-resolution instruments with advanced ion collection methods ^1,16-18^. Recently, two studies employed comparable TMT10-plex strategies to determine the expression level of proteins in four human cell lines ^19^ and ovarian tumour sections ^20^. Both studies used high pH RP to fractionate the pooled TMT-labeled peptide sample into 96 fractions, which were concatenated into 10 and 12 samples, respectively. These samples were subsequently subjected to low pH RP on an EASY-nLC 1000 LC pump coupled to an Orbitrap Fusion mass spectrometer with MultiNotch MS3 analysis ^17,19,20^. In total, 8590 proteins were quantified across 10 human cell lines in 36 h of LC-MS time ^19^ and 8167 proteins were quantified across 18 ovarian tumour samples in ~24 h ^20^. This proteome coverage is consistent with the depth observed by Spicer *et al* (8757 proteins) using a similar concatenated 2D RP-RP LC-MS/MS method with non-labeled samples ^4^.

Notably, Bekker-Jensen *et al* improved coverage of the human proteome to 10284 protein-coding genes from tryptic digests of HeLa cells using a 2D RP-RP LC-MS/MS approach without concatenation of 46 fractions by increasing peptide loads, optimizing online LC-MS gradients, and employing a fast scanning Q-Exactive HF instrument ^10^. The acquisition required only 34.5 h of LC-MS/MS time, and the authors suggest that running short gradients on many fractions may provide a suitable compromise between instrument time used and proteome coverage obtained ^10^. A new LC system from Evosep Biosystems (Denmark) appears to expand on this concept. The Evosep One LC system elutes peptides from a special C18 StageTip, called the Evotip, at low pressure and then captures the gradient and eluted analytes in a long storage loop ^21^. A single high-pressure pump then applies the stored gradient with embedded, pre-separated peptides to a C18 analytical column. The Evosep system has pre-set methods optimized to provide the best performance to time compromise based on sample complexity, with gradient lengths of 21 or 44 min for more complex samples ^21^. Building on the workflow strategy of Bekker-Jensen *et al* ^10^, Bache *et al* performed a comparative analysis of 46 HeLa fractions by Evosep One with the 60 samples per day method (21 min gradients) versus 15 min gradients on an EASY-nLC 1200. The Evosep One method identified 9918 proteins in 18.4 h, while the EASY-nLC method identified 9603 proteins in 28.3 h, indicating that Evosep One technology can significantly reduce measurement time without diminishing performance ^21^.

As part of a larger effort to attain quantitative proteomics data on >100 primary nonsmall cell lung carcinoma xenografts ^22^, we wished to determine if the Evosep One platform could be incorporated into our multiple-dimension LC-MS/MS and extended TMT multiplexing workflow. We TMT-labeled xenograft samples, pooled and separated the samples by high pH RP into 60 fractions, and then analyzed the fractions by a concatenated EASY-nLC conventional workflow, 30 samples per day Evosep workflow, 60 samples per day Evosep workflow, and concatenated Evosep workflow. We show that Evosep technology can be used for the simultaneous identification and quantification of complex TMT-labeled protein mixtures in less time without compromising depth of proteome coverage.

## Experimental Procedures

### Sample preparation and TMT labeling

Samples of nonsmall cell lung carcinoma (NSCLC) patient-derived xenografts (PDX) were obtained as previously described ^22,23^. Tumour samples were wetted with cold PBS, weighed, and mixed with lysis buffer (0.5 M Tris pH 8.0, 50 mM NaCl, 2% SDS, 1% NP-40, 1% Triton X-100, 40 mM chloroacetamide, 10 mM TCEP, 5 mM EDTA) in a ratio of 10 mg wet tissue to 200 μL buffer. Tumours were cut into small pieces, sonicated for 15 sec twice, heated at 95°C for 20 min with mixing at 1000 rpm, cooled to room temperature (~10 min on bench) and centrifuged at 20000 g for 5 min at 18°C ^20^. Protein concentration of the supernatant was measured by tryptophan fluorescence assay ^24^. 100 μg of proteins were precipitated by methanol-chloroform method and digested overnight at 37°C with 1 μg trypsin/Lys-C mixture (Promega Cat#V5073) ^19^. Peptide concentration was measured by A_280_ and 50 μg labeled with 10-plex tandem mass tag reagents according to manufacturer’s instructions (Thermo Fisher Scientific, Cat#90110). Eight PDX samples were grouped and individually labeled with isobaric compounds (TMT10-126, 127, 128N, 129, or 130), while two pooled mixtures of all eight PDX samples were labeled with TMT10-128C or 131, to serve as normalization controls ^16,20^. Excess TMT label was quenched with 8% ammonium hydroxide prior to TMT-labeled samples being pooled at a 1:1:1:1:1:1:1:1:1:1 ratio, vacuum centrifuged, and stored at −80°C.

### High-pH reversed-phase fractionation

Lyophilized TMT mixes were resuspended in 20 μL of ddH_2_0 and fractionated using high pH reversed phase high-pressure liquid chromatography (HPLC) at 4°C. We used a Waters 1525 binary HPLC pump and Waters XBridge Peptide BEH C18 4.6mm × 250mm column (Part 186003625, Waters, USA) with a flow rate of 1mL/min. A 90 min gradient using buffer A (ddH_2_O, adjusted to pH 10 with ammonium hydroxide) and buffer B (80% Acetonitrile, adjusted to pH 10 with ammonium hydroxide) was run as follows: 0-3 min 1-12% B; 3-60 min 12-30% B; 60-65 min 30-60% B; 65-70 min 60-99% B, 70-75 min 99-1% B, 75-90 min 1% B. Ultra violet (UV) absorbance was measured throughout the gradient at 214 nm and 280 nm using a Waters 2489 UV/Visible detector. Fractions were collected from the beginning of the gradient in 1.2 minute time intervals for 60 fractions (72 min). The remainder of the gradient was not collected as this served as column cleaning and re-equilibration.

### LC-MS/MS analysis

For the traditional LC-MS/MS workflow, the 60 fractions obtained from high pH separation were lyophilized, resuspended in 100 μL of 0.1% formic acid, and concatenated into 15 samples by combining multiple early, middle, and late fractions (eg. 1+16+31+46). Samples were lyophilized, resuspended in 80 μl 0.1% TFA, and de-salted on homemade C18 stagetips. Subsequently, samples were analyzed by LC-MS using an EASY-nanoLC 1200 system with a 4-hour gradient and an Orbitrap Fusion Lumos Tribrid mass spectrometer (Thermo Fisher Scientific, USA). For the Evosep workflow, the 60 high pH fractions were lyophilized, resuspended in 100 μL of 0.1% formic acid, and transferred to a 96-well plate. Each fraction was loaded in its entirety onto a disposable Evotip C18 trap column (Evosep Biosytems, Denmark) as per manufacturer’s instructions. Briefly, Evotips were wetted with 2-propanol, equilibrated with 0.1% formic acid, and then loaded using centrifugal force at 1200 g. Evotips were subsequently washed with 0.1% formic acid and then 200 μL of 0.1% formic acid was added to each tip to prevent drying. Samples were injected into the same Orbitrap Fusion Lumos MS as above using an Evosep One instrument (Evosep Biosystems). Standard pre-set methods were used (60 or 30 samples per day) for the LC component of the run. For the Evosep concatenated workflow, the 60 fractions were concatenated into 30 samples (1+30, 2+31…29+60) and loaded with Evotips using the 60 samples per day method. For all workflows, peptides were introduced by nano-electrospray into the mass spectrometer using an Easy-Spray source (Thermo Fisher Scientific) and data were acquired using MultiNotch synchronous precursor selection (SPS) MS3 scanning for TMT tags ^15^. Overall, monoisotopic precursor selection (MIPS) was determined at the peptide level, with a global intensity threshold of 1 × 10^4^, and only peptides with charge states of two to six were accepted. MS1 acquisition resolution was set to 120,000 with an automatic gain control (AGC) target value of 4 × 10^5^ and maximum ion injection time (IT) of 50 ms for a scan range of 550-1800 m/z. Isolation for MS2 scans was performed in the quadrupole, with an isolation width of 2 m/z. MS2 scans were performed in the linear ion trap with maximum ion IT of 50 ms, AGC target value of 1 × 10^4^, and normalized collisional energy (NCE) of 35 using the turbo scan rate. For MS3 scans, ions were isolated in the quadrupole with an isolation width of 2 m/z. Higher-energy collisional dissociation (HCD) activation was employed, with NCE of 65, and scans were measured in the orbitrap with resolution of 50,000, scan range of 100-500m/z, AGC target value of 1 × 10^5^, and maximum ion IT of 50 ms. Dynamic exclusion was set to 65 s and 45 s for the 30 samples per day and 60 samples per day methods, respectively, to avoid repeated sequencing of identical peptides.

## Data analysis

MS raw files were analyzed using Proteome Discoverer 2.2 (Thermo Fisher Scientific) and fragment lists searched against the human and mouse UniProt Reference databases (Human downloaded Jan 15, 2018, 48177 entries, reviewed entries only; Mouse downloaded Oct 25 2017, 25074 entries, reviewed entries only) by both Sequest HT and MS Amanda 2.0 search engines ^25^. For both search algorithms, the parent and fragment mass tolerances were set to 10 ppm and 0.6 Da, respectively. Only complete tryptic peptides with a maximum of two missed cleavages were accepted. Methionine oxidation, protein N-terminal acetylation, and asparagine and glutamine deamidation were included as variable modifications, while cysteine carbamidomethylation and TMT labelling of peptide N-termini and lysine residues were considered fixed modifications. Search results were filtered through Percolator ^25^ at the peptide spectral match (PSM) level using a strict false discovery rate (FDR) *q*-value of 0.01 and relaxed FDR *q*-value 0.05 ^26^. TMT reporter ions were quantified using the Proteome Discoverer 2.2 reporter ions quantifier node with an integration tolerance of 20 ppm, on the MS order of MS3. The standard consensus workflow for reporter ion quantification was used. The mass spectrometry proteomics data have been deposited to the ProteomeXchange Consortium via the PRIDE partner repository with the dataset identifier PXD012520 and 10.6019/PXD012520.

## Results and Discussion

To perform quantitative whole proteome analysis of NSCLC PDX samples in a multiplexed manner, eight PDX samples and two reference samples (a mixture of all PDX samples) were digested and labeled with isobaric tags. Labeled peptides were combined together and then separated into 60 fractions using high pH reversed phase HPLC fractionation, dried under vacuum centrifugation, resuspended and concatenated into 15 final samples, and de-salted on homemade C18 stagetips. Subsequently, samples were analyzed by LC-MS using an EASY-nLC 1200 system with a 4-hour gradient and an Orbitrap Fusion Lumos Tribrid mass spectrometer operating in MS3 mode. Under these experimental conditions, 12789 proteins were identified, 9359 with greater than two unique peptides (Table 1, Condition A). This depth of coverage is consistent with that observed for human cell lines ^19^. Increasing the amount of starting material from 0.5 mg to 1 mg did not result in more protein identifications (Table 1, Condition B). Indeed, fewer proteins were identified when more starting material was used, suggesting that 0.5 mg of starting material is sufficient for optimizing protein identification by this method. Assuming an average LC equilibration and load time of 30 minutes, this conventional workflow consumed 67.5 hours of mass spectrometry time, which translates to 3.2 protein identifications per minute of MS time (Figure 1 and Table 1, Condition A).

**Table 1.** Comparison of proteome coverage for different experimental workflows. The number of proteins and peptides identified for experimental conditions A) Easy-1200 concatenated, 0.5 mg starting material, B) Easy-1200 concatenated, 1 mg starting material, C) Evosep One, 30 samples per day method, D) Evosep One, 60 samples per day method (three replicates and their average are shown) and E) Evosep One concatenated, 60 samples per day method. All Evosep One conditions employed 0.5 mg of starting material, and entire fractions were loaded.

| Condition | Experimental Description | Number of proteins identified, 1 unique peptide | Number of proteins identified, $\geq 2$ unique peptides | Number of peptides | Hours of mass analysis time | Proteins identified per MS minute |
| --- | --- | --- | --- | --- | --- | --- |
| A | EASY-nLC concatenated, 0.5 mg starting material | 12789 | 9359 | 87554 | 67.5 | 3.2 |
| B | EASY-nLC concatenated, 1 mg starting material | 11731 | 7721 | 68806 | 67.5 | 2.9 |
| C | Evosep One, 30 samples per day, 0.5 mg starting material | 12339 | 9008 | 79888 | 48 | 4.3 |
| D (1) | Evosep One, 60 samples per day, 0.5 mg starting material | 10212 | 8106 | 57460 | 24 | 7.1 |
| D (2) | Evosep One, 60 samples per day, 0.5 mg starting material | 10825 | 8417 | 73690 | 24 | 7.5 |
| D (3) | Evosep One, 60 samples per day, 0.5 mg starting material | 10419 | 8220 | 56549 | 24 | 7.2 |
| D (Avg) | Evosep One, 60 samples per day, 0.5 mg starting material | 10485 | 8248 | 62566 | 24 | 7.3 |
| E | Evosep One Concatenated, 30 samples per day, 0.5 mg starting material | 8137 | 5468 | 48209 | 12 | 11.3 |

**Figure 1.**
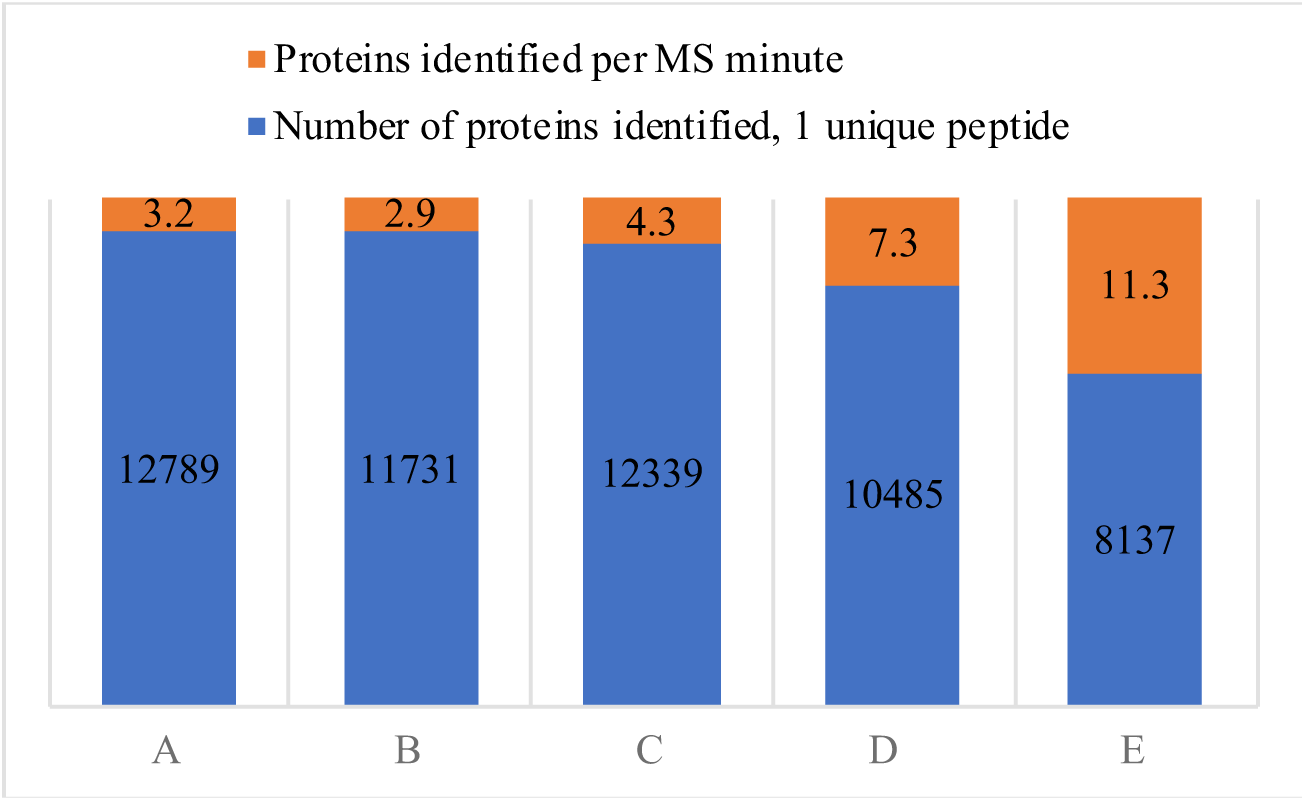
Comparison of proteome coverage and efficiency of coverage for traditional versus Evosep workflows. The number of proteins identified (≥ one unique peptide) and number of proteins identified per minute of mass analysis time are shown in blue and orange, respectively, for conditions A) Easy-1200 concatenated, 0.5 mg starting material, B) Easy-1200 concatenated, 1 mg starting material, C) Evosep One, 30 samples per day method, D) Evosep One, 60 samples per day method (numbers represent average of three replicates), and E) Evosep One concatenated, 60 samples per day method.

To determine if the new Evosep One LC system could achieve similar numbers of protein identifications for multiplexed samples while using less mass spectrometry time, we established a novel workflow for the same TMT labelled PDX samples involving Evotips. High pH reversed phase HPLC fractionated 0.5 mg of starting material into 60 fractions, which were then lyophilized, resuspended in 100 μl of 0.1% formic acid and loaded onto an Evotip. Given that Evotips have capacity to bind ~ 1-2 μg of material, this strategy of overloading the Evotips essentially regulates the load and prevents overloading of the analytical column. Peptides were eluted from the Evotip using a 44 min gradient and 15 cm analytical column (100 μm i.d., 3 μm C18 beads) with the Evosep One LC system and 30 samples per day method. Peptides were mass analyzed on the same Orbitrap Fusion Lumos instrument as used in the concatenated EASY-nLC workflow, operating in MS3 mode. We found that the number of proteins identified with at least one unique peptide (12339) or two unique peptides (9008) was comparable to the EASY-nLC workflow (Table 1, Condition C). However, the new Evosep workflow used only 48 hours of mass spectrometry time, a 29% reduction in measurement time from the EASY-nLC workflow, resulting in 4.3 protein identifications per minute of MS time (Table 1, Condition C). This indicates that the use of Evotips and Evosep One in the mass analysis of multiplexed samples can significantly decrease mass spectrometry time without compromising data quality.

Given our initial success with the new Evosep workflow, we wished to ascertain how shortening the Evotip elution gradient would impact depth of proteome coverage. The Evosep workflow was repeated with a 21 min elution gradient and 8 cm analytical column (100 μm i.d., 3 μm C18 beads) on the same TMT labeled PDX samples. This 60 samples per day workflow resulted in the identification of 10212 proteins with at least one unique peptide and 8106 with at least two unique peptides (Table 1, Condition D(1)). Similar depth of proteome coverage was observed for biological repeats (Table 1, Conditions D(2) and (3)), indicating that our new Evosep workflow for TMT labeled samples is reproducible. Indeed, 84.1% of proteins identified with two unique peptides were identified in all three replicates, with 97.2% identified in at least two of the three replicates (Figure 2A). With three unique peptides, 93.7% of proteins were identified in all three replicates, and 99.5% of proteins identified in at least two replicates (Figure 2B). As well, the number of MS/MS spectra and peptide spectral matches (PSMs) observed per fraction was consistent between replicates (Figure 2C). On average, the 60 samples per day Evosep workflow resulted in 13% and 10% less protein identifications (two peptides) compared to the traditional and 30 samples per day workflows, respectively. However, the 60 samples per day workflow also reduced mass spectrometry time by 64% and 50% compared to traditional and 30 samples workflows, respectively, resulting in 7.3 protein identifications per minute of MS time on average (Figure 1, Condition D). This suggests that in situations where mass spectrometry time is limited or cost prohibitive, it is feasible to mass analyze 60 multiplexed samples per day with the Evosep One without drastically reducing proteome coverage. Indeed, 76% of identified proteins were identified in all three workflows (Figure 2D, 8092/10616). Furthermore, gene ontology analysis revealed consistent distribution of proteins across three ontology classifications for the concatenated EASY-nLC workflow and the 30 samples per day Evosep workflow (Figure 3). It should be noted that in this particular study, the number of identified proteins is inflated since we are using a two-species xenograft system, such that the increase in protein identification by the concatenated EASY-nLC workflow compared to Evosep workflows may not be as substantial with a single species model.

**Figure 2.**
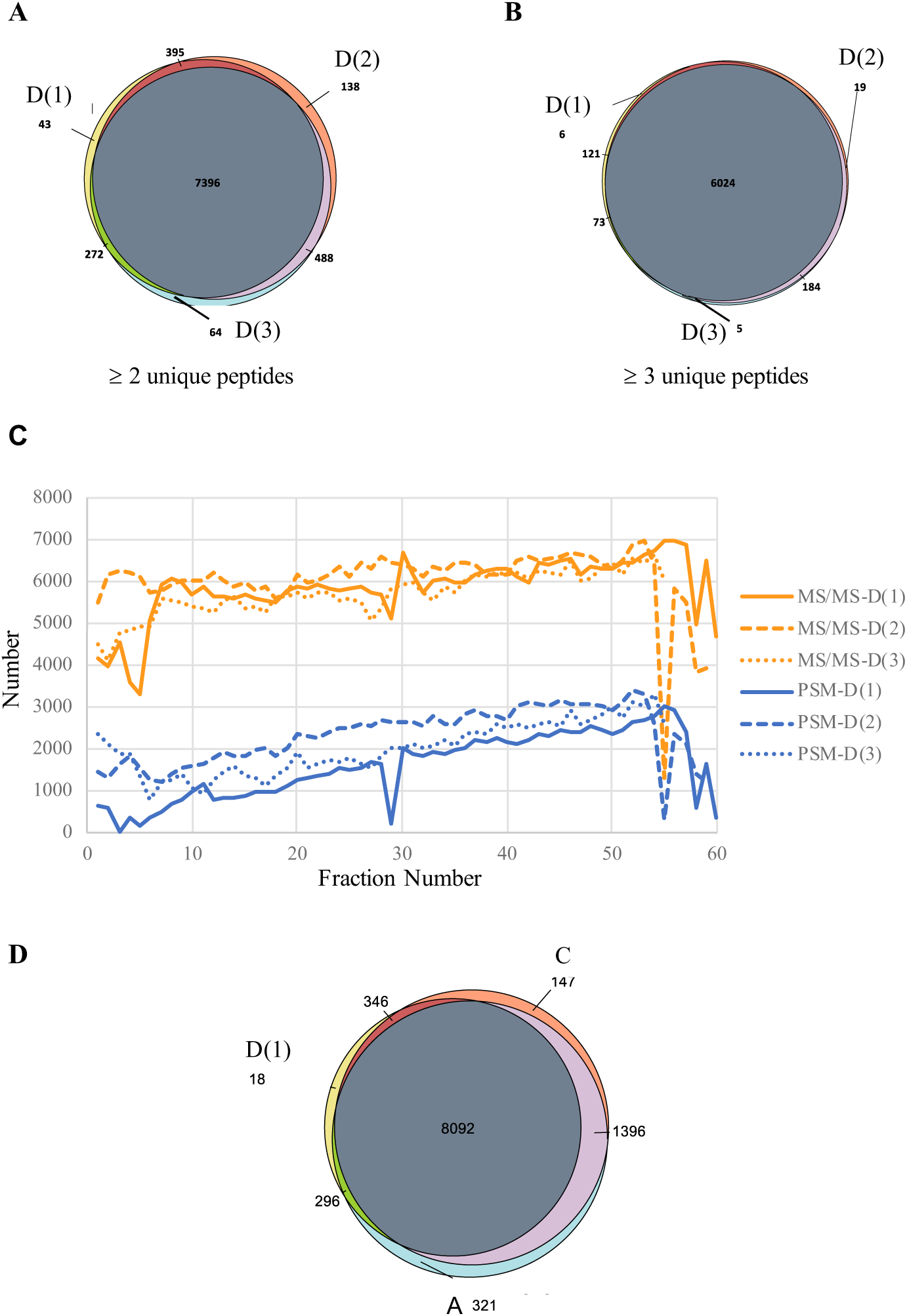
Reproducibility of Evosep replicates and consistency of coverage between Evosep and traditional LC-MS/MS workflows. Total protein identifications with A) ≥ two unique peptides or B) ≥ three unique peptides are shown in Venn diagrams comparing three replicates (D(1), D(2), and D(3)) of the Evosep 60 samples per day workflow. C) The number of MS/MS spectra and PSMs for the Evosep replicates are shown in orange and blue, respectively. Total protein identifications with D) ≥ two unique peptides are shown in Venn diagrams comparing concatenated LC-MS/MS and Evosep workflows.

**Figure 3.**
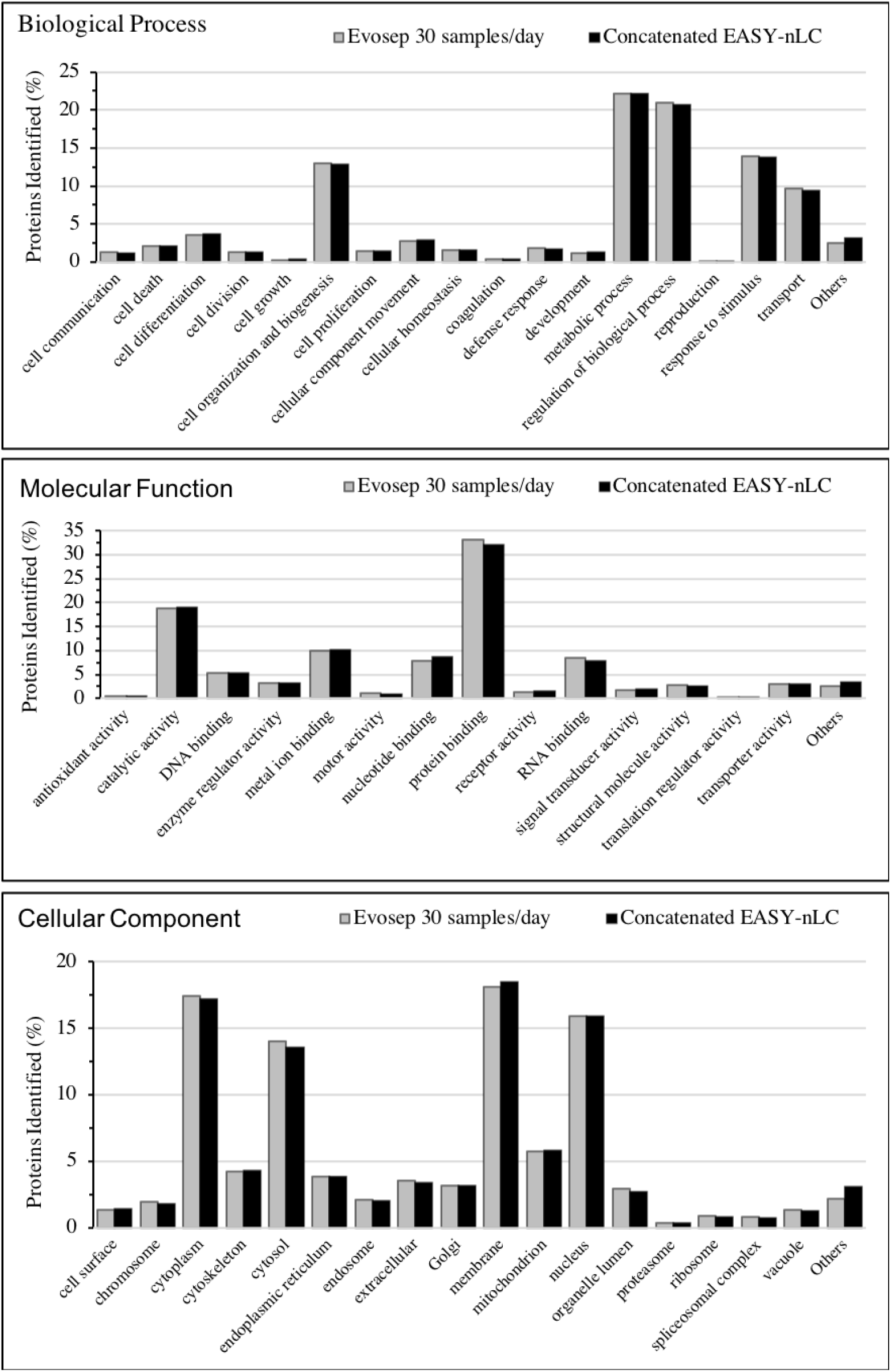
Gene ontology analysis. Analysis of biological process, molecular function, and cellular component was performed in Proteome Discoverer 2.2. The percentage of proteins identified in each gene ontology classification is shown for the 30 samples per day Evosep and EASY-nLC workflows.

Our TMT 10-plex sample group consisted of eight PDX samples and two pools of the eight samples that served as normalizing mixtures. The normalized relative abundance of proteins identified in each isobaric channel in each replicate was found to be consistent across all samples processed using Evosep workflows (Figure 4A). Given that the purpose of including two control samples within each sample group was to allow normalization across groups, it is important that a large number of the same proteins are identified in each control sample replicate. Thus, we analyzed the proteins identified in the common pooled samples (labelled with 128C and 131) across the three Evosep biological replicates. An average of 64.2% of all identified proteins were found in all three pooled control replicates (Figure 4B, C), suggesting that this workflow will provide sufficiently large numbers of proteins for normalization between groups. To assess intra-group variability, we compared proteins identified in the control sample labelled with 128C to the identical sample labelled with 131. With a low threshold of one unique peptide, we found that less than 0.3% (+/-0.1%) of total proteins from the mixed sample were identified in only one of the two control samples, indicating consistent labelling of samples. Notably, 79.9% of all PSMs acquired via the Evosep workflows had a Percolator ^25^ q-value <0.001, and 96.3% of all PSMs had a q-value <0.005, indicating peptides are high quality.

**Figure 4.**
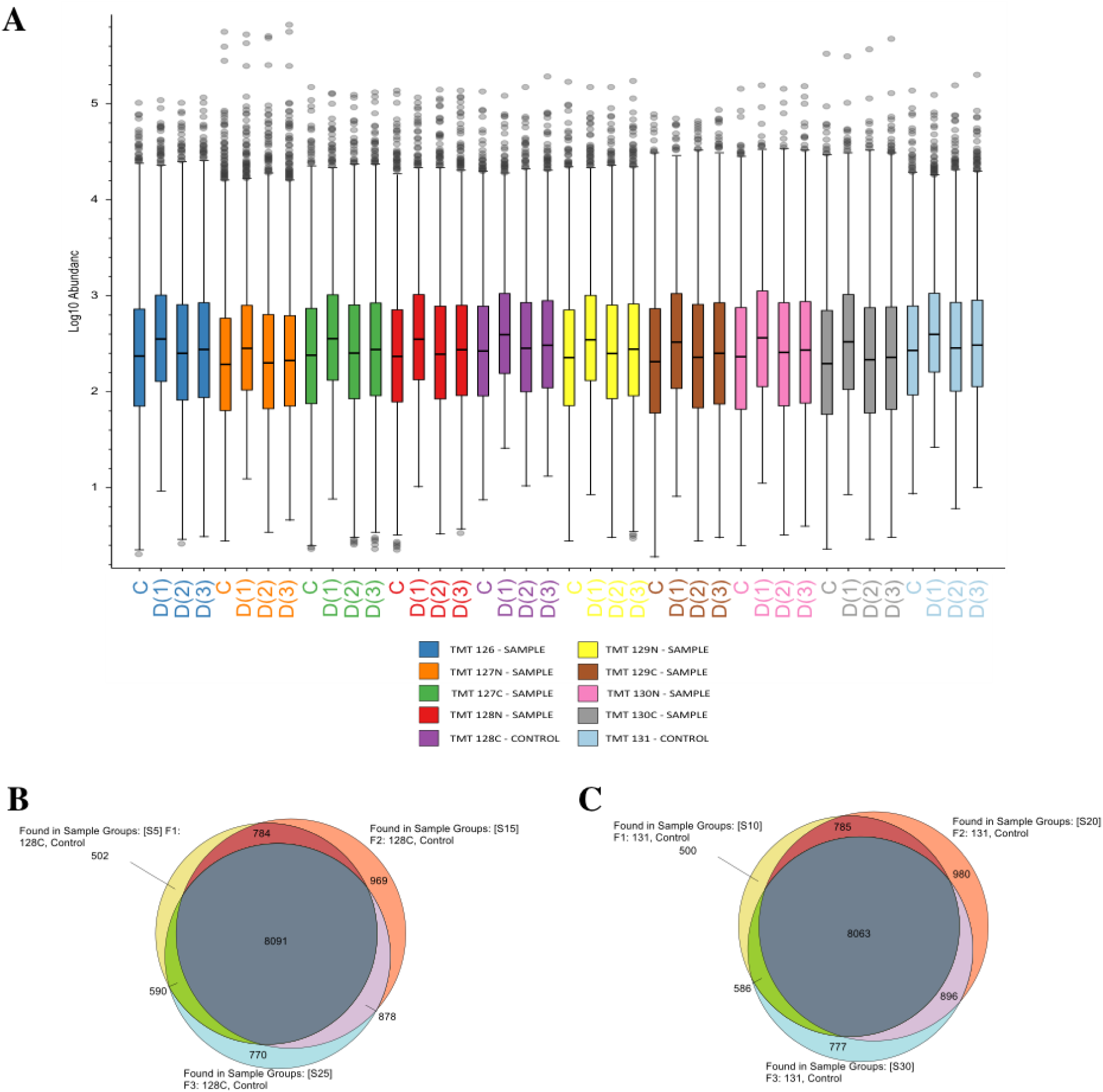
Relative protein abundance and protein identifications across Evosep replicates. A) The Log10 normalized abundance of proteins by channel is shown for four Evosep workflow experiments. F4 represents the 30 samples per day Evosep experiment, while F1, F2, and F3 represent the 60 samples per day Evosep replicates. In all cases, samples were independently labelled. Total protein identifications with ≥ 2 unique peptides are shown in Venn diagrams comparing B) TMT-128C and C) TMT-131 control groups between 60 samples per day Evosep replicates.

A key aspect to deep proteome analysis is the offline peptide separation strategy employed. Ideally, this strategy separates peptides in a manner that minimizes overlap between adjacent fractions. To assess the effectiveness of our high pH reversed phase separation in the first dimension, we examined peptides that were common between fractions. We found an average 11.8% (±1.1%) peptide overlap in adjacent side by side fractions across all fractions and all replicates (Figure 5A). Peptide overlap in alternate (separated by two) and distant (separated by five) fractions reduced to 3.9% (±0.9%) and 1.5%, (± 0.4%), respectively (Figure 5B, C). This indicates that our first-dimension separation strategy efficiently and effectively separates complex PDX samples. As well, the number of MS/MS spectra observed per fraction was relatively uniformly distributed across the 60 fractions (Figure 2C), consistent with good orthogonality of our high pH -low pH HPLC/Evotip separation protocol. We noticed that very early and very late fractions (1-5 and 55-60, Figure 2C) showed a marked reduction of MS/MS spectra, suggesting that these fractions may be combined with middle fractions to further reduce mass analysis time. To test the impact of concatenation of fractions prior to Evotip processing, we combined the 60 HPLC fractions into 30 fractions and proceeded with the Evosep 60 samples per day workflow as described above. While this experimental design reduced the mass analysis time to 12 hours, it also resulted in significantly fewer protein identifications (Table 1). Specifically, the concatenated Evosep strategy produced 22.4% and 34.1% less protein identifications than when the 60 fractions were run individually with 60 samples per day and 30 samples per day workflows, respectively (Table 1 and Figure 1). Thus, unless mass spectrometry time is extremely limited or cost prohibitive, it is recommended that HPLC fractions not be concatenated prior to the Evosep One workflow. However, concatenating just the very early and late fractions may not negatively impact proteome coverage.

**Figure 5.**
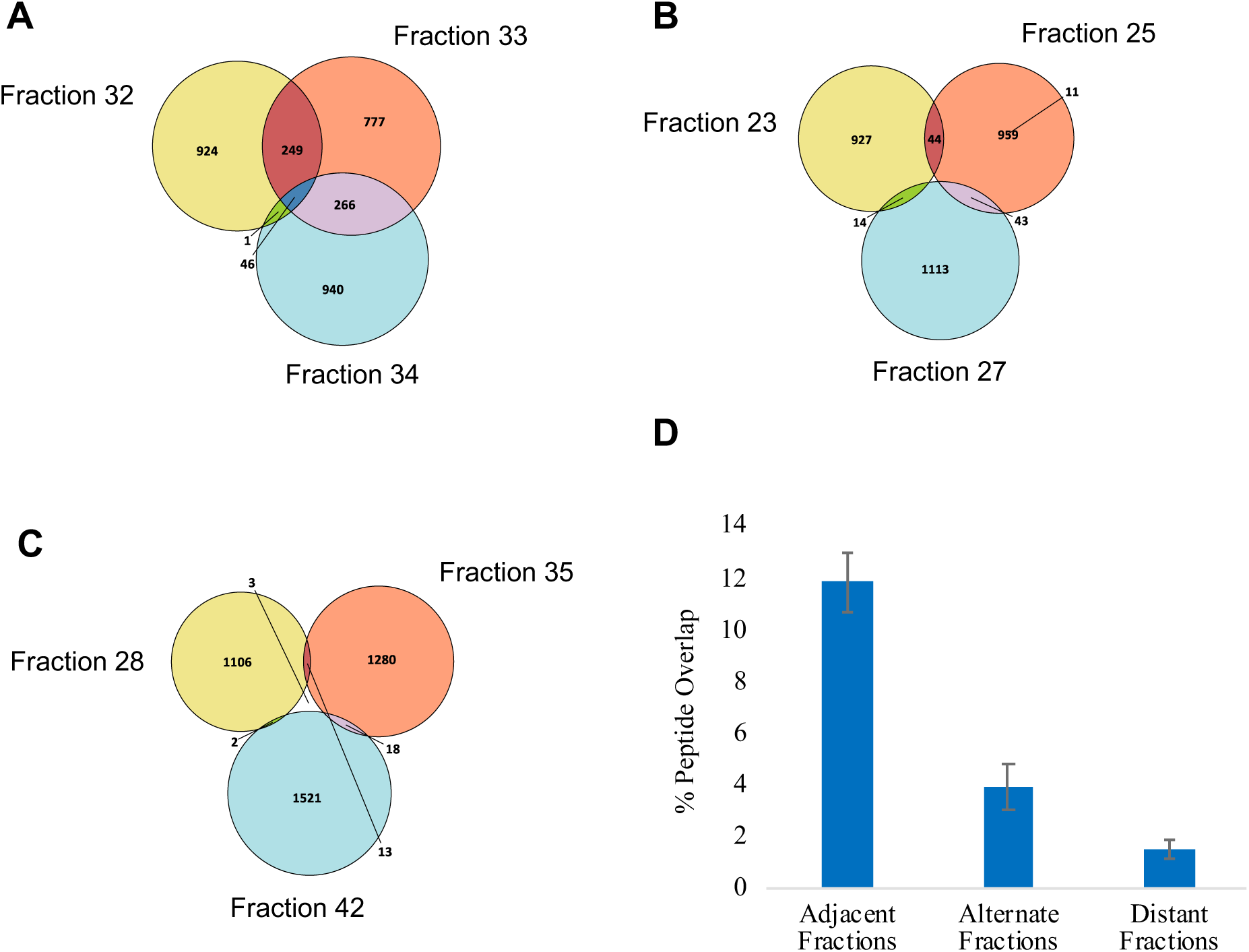
Separation of peptides high pH reversed phase fractionation. Venn diagrams show peptides observed in representative A) adjacent, B) alternate, and C) distant fractions from the same replicate. D) The average percentage of peptide overlap for each category of fraction is shown for the three Evosep replicates, with error bars representing standard error.

## Conclusions

Achieving maximal efficiency of MS time and deep quantifiable proteome coverage is a collective goal in the field of proteomics. As the demand for deep proteomic analyses of cell lines and tissues increases, it is imperative that workflows optimize proteome coverage in a cost and time efficient manner. While the Evosep One liquid chromatography system was constructed for high throughput applications with a particular focus on clinical analysis ^21^, we show that this technology can be used to support data dependant acquisition of unknown TMT labelled samples. Incorporating Evosep One into our TMT workflow reduced the MS analysis time drastically while maintaining data quality. The 30 samples per day method (total MS time of 48 hours) gave the best compromise between deep proteome coverage and MS analysis time. Each TMT-labelled group contained eight unknown NSCLC PDX samples, such that our Evosep workflow achieved deep proteome coverage in only six hours per sample, making deep proteome analysis of large sample numbers more manageable. The 60 sample per day method as well as the concatenated 60 sample per day method also provided reasonable proteomic depth while significantly reducing instrument time. Nevertheless, the cooperation between the first dimension of separation (gradient length and number of fractions) and Evosep throughput (gradient length) has not been fully explored, creating opportunity to further optimize proteome coverage versus mass analysis time. This will become especially evident as mass spectrometry increases in acquisition speed and sensitivity.

## AUTHOR INFORMATION

## Author Contributions

The manuscript was written through contributions of all authors. All authors have given approval to the final version of the manuscript.

## Notes

As an employee of Evosep, NB has a potential conflict of interest regarding this work. The other authors declare no competing financial interest.

### Abbreviations

2D-LC: two-dimensional liquid chromatography
MS: mass spectrometry
NSCLC: non-small cell lung carcinoma
PDX: patient-derived xenografts
RP: reversed phase
TMT: tandem mass tag

